# The effects of adolescent stress on adult social behavior and basolateral amygdala GABAergic neurons with perineuronal nets depend on prenatal stress history

**DOI:** 10.1101/2025.07.31.667823

**Authors:** Marcia C. Chavez, Madeline M. Jones, Alyssa R. Whaley, Trevor M. Pounders, Jessica T. Tremblay, Megan Zajkowski, Maria Ragusa, Billy Y. Lau, Kalynn M. Schulz

## Abstract

Developmental stress is a well-established risk factor for mental health disorders, yet the neural mechanisms underlying these outcomes remain incompletely understood. Inhibitory brain networks, particularly within the amygdala, are disrupted by stress and implicated in stress-related psychopathologies. Using a rodent model, the current study investigated the isolated and combined effects of prenatal and adolescent stress on adult social interactions and GABAergic neurons surrounded by perineuronal nets (PNNs) in the basolateral amygdala (BLA). Male and female rats were exposed to chronic variable stressors (CVS) prenatally (PS), during adolescence (AS), or during both prenatal and adolescent periods (PS+AS). In adulthood, all animals were tested for social behavior with same-sex weight-matched partners, and brains were collected for identification of BLA inhibitory neurons (GAD67 staining) and PNNs (*Wisteria Floribunda Agglutinin* staining). For social behavior, AS alone robustly increased social investigation in adulthood relative to non-stressed (NS) controls and animals exposed to combined PS+AS. PS+AS subjects did not significantly differ from NS controls, suggesting that prenatal stress exposure prevented adolescent stress-induced increases in adult social investigation. An analogous data pattern was observed in the BLA. AS alone decreased the number GAD67+ neurons surrounded by PNNs (co-labeled) relative to NS controls and subjects exposed to combined PS+AS. When the percentage of total GAD67+ neurons co-labeled with PNNs was assessed, both PS alone and AS alone reduced the proportion of GAD67+ neurons surrounded by PNNs, whereas combined PS+AS had no effect. Overall, these data suggest that prenatal stress exposure prevents adolescent stress-induced disruptions to perineuronal nets surrounding inhibitory neurons in the BLA, potentially conferring resilience to adolescent stress-induced changes in inhibitory function and social behavior.

**Highlights:**

- Adolescent stress exposure increased social investigation in adulthood.
- Adolescent stress decreased the number of BLA cells co-labeled with GAD67 and WFA.
- Prenatal or adolescent stress decreased the proportion of inhibitory neurons with PNNs.
- When preceded by prenatal stress, effects of adolescent stress were not observed.

---

1. **Introduction**

Brain inhibitory networks are dysregulated by stress exposure (Busler et al., 2022; Jie et al., 2018) and also implicated in mental health disorders with etiologic links to stress experienced during development (Duman et al., 2019; Nuss, 2015). For example, amygdala activity is elevated in individuals diagnosed with stress-related psychopathologies (Groenewold et al., 2013), which may be due to compromised inhibitory function (Babaev et al., 2018; Nuss, 2015). The basolateral amygdala (BLA) is a key region within the neural network mediating stress responsivity and the expression of anxiety-related and social behavior (Hetzel & Rosenkranz, 2014; Lalumiere, 2014; Rosenkranz et al., 2010). In adult rodents, decreased inhibition and/or increased excitation of the BLA decreases social investigation (Asim et al., 2024; Paine et al., 2017; Rosenkranz et al., 2010; Sanders & Shekhar, 1995; Truitt et al., 2007), and previous studies indicate that stress-induced BLA excitation is the consequence of decreased inhibitory regulation (Asim et al., 2024; Truitt et al., 2007). Importantly, while the effects of stress experienced in adulthood generally diminish over time (Tran & Gellner, 2023), stress encountered during prenatal or adolescent development results in long-lasting or even permanent changes in behavior. Rodents exposed to either prenatal or adolescent stress exhibit increased anxiety-like behaviors in adulthood (P. R. Lee et al., 2007; Schulz et al., 2011, 2014), accompanied by reductions in markers of inhibitory function within the amygdala (Albrecht et al., 2017; Tzanoulinou et al., 2014; Zhu et al., 2018). The current study aims to extend previous findings by investigating the long-term impacts of combined prenatal and adolescent stress exposure on social behavior and inhibitory neurons within the BLA.

The function of inhibitory neurons is influenced by perineuronal nets (PNNs). PNNs are stress-sensitive extracellular matrix structures that preferentially surround inhibitory neurons and their synaptic contacts (Laham & Gould, 2022). Several studies demonstrate that PNNs regulate and enhance inhibitory neuron function by limiting the formation of new synaptic contacts and strengthening existing synaptic connections (Bosiacki et al., 2019; Favuzzi et al., 2017). In adulthood, enzymatic removal of PNNs can reduce network-level inhibition within the hippocampus (Liu et al., 2023), and also cortical areas such as the medial prefrontal cortex (Carceller et al., 2020), entorhinal cortex (Christensen et al., 2021) and visual cortex (Christensen et al., 2021; Lensjø, Lepperød, et al., 2017; Liu et al., 2023). To date, only a handful of studies have investigated the long-term impacts of adolescent stress on PNNs (Colodete et al., 2024; Page & Coutellier, 2018; Santos-Silva et al., 2024). In the short-term, adolescent corticosterone exposure decreases the percentage of BLA inhibitory neurons surrounded by PNNs (Virakorn et al., 2024), however, whether these effects of adolescent stress persist into adulthood is currently unknown. Likewise, although prenatal stress disrupts aspects of GABAergic neuron development (Lussier & Stevens, 2016; Stevens et al., 2013), whether prenatal stress impacts adult levels of PNNs surrounding inhibitory neurons has not been investigated in any brain region, nor have the combined effects of prenatal and adolescent stress exposures. Therefore, we sought to examine the long-term impact of prenatal, adolescent, and combined prenatal and adolescent stress on GABAergic neurons and PNNs in the BLA.

Individuals experience multiple stressors throughout life, yet the consequences of combined stress exposure during both prenatal and adolescent periods has received little investigation. In contrast, the cumulative health risks of postnatal stressors are well documented in humans (Björkenstam et al., 2015; Evans et al., 2013; Felitti et al., 1998; Hanson et al., 2016), consistent with the concept of allostatic load resulting from the accumulation of stressors over time (Gallo et al., 2014; McEwen & Stellar, 1993). Stressors may have unique effects when experienced across pre- to post-natal development. According to the predictive adaptive response model (Gluckman et al., 2005; Gluckman & Hanson, 2004), prenatally stressed animals may have an adaptive advantage if similar conditions are encountered postnatally. Here we sought to distinguish whether the effects of prenatal and adolescent stress on adult social behavior and GABAergic neurons are cumulative, or whether prenatal stress changes the impact of adolescent stress exposure. Our findings align with the predictive adaptive response model, and we report here that prenatal stress exposure prevents adolescent stress-induced alterations in social behavior and GABAergic neurons surrounded by PNNs in the basolateral amygdala.

## 2. Materials and Methods

### 2.1 Animals

Animals were maintained on a 12:12 light/dark cycle with lights on at 07:00 and a vivarium temperature of 21°C. All animals had ad libitum access to food (2018 Teklad Global 18% Protein RodentDiet, Harlan Laboratories Inc., Indianapolis, IN) and water. Sixty male and female Sprague Dawley rats were paired to generate experimental animals. The day a vaginal plug was observed was designated as gestation day 0. Following parturition, food and water continued to be replaced weekly, but the bedding and nests were left undisturbed until weaning at 21 days of age to minimize stress. Cage cleanliness was closely monitored during this time, and additional bedding was provided if necessary (Tekfresh, Harlan Laboratories Inc., Indianapolis, IN). Upon weaning, weekly cage changes resumed, and animals were housed 2 per cage with same-sex littermates. Two to four male and female offspring were weaned from a litter and housed with a same sex, and same stress condition littermate. A total of 138 animals were used in the current study (70 male, 68 female). Animals were treated in accordance with the NIH Guide for the Care and Use of Laboratory Animals, and all protocols were approved by the Denver VAMC Animal Care and Use Committee.

### 2.2 Experimental design and stress procedures

A 2x2x2 between-subjects research design was employed to investigate the effects of prenatal condition (prenatal stress vs. no prenatal stress), adolescent condition (adolescent stress vs. no adolescent stress), and sex (male vs. female) on adult social behavior and PNNs surrounding GABAergic neurons in the BLA complex (Figure 1A). This fully factorial design generated four groups: prenatal stress (PS, n = 32), adolescent stress (AS, n =32), combined prenatal and adolescent stress (PS+AS, n = 32), and non-stressed (NS, n = 42). Each experimental group had an equal number of males (n = 16) and females (n = 16) with the exception of the NS group that had 22 males and 20 females. Detailed descriptions of our chronic variable stress (CVS) procedures have been published previously (Chavez et al., 2021; Schulz et al., 2011, 2013). In brief, pregnant rat dams were randomly assigned to receive either chronic variable stressors 1-4 times daily during gestation days 14-21, or were left undisturbed during gestation. Chronic variable stressors included body restraint in a plastic cylinder tube (60 minutes), transportation on a noisy cart around the animal facility (30 minutes), forced swim in water between 22-25 °C (5 minutes), social crowding (6-12 same-sex rats per cage, 8 hours), overnight fast (15 hours), continuous light (36-48 hours), and cold room exposure (6 hours, 4°C). Offspring randomly assigned to the adolescent stress group underwent these same stress procedures from postnatal day 23 to 51 (Figure 1B).

**Figure 1.**
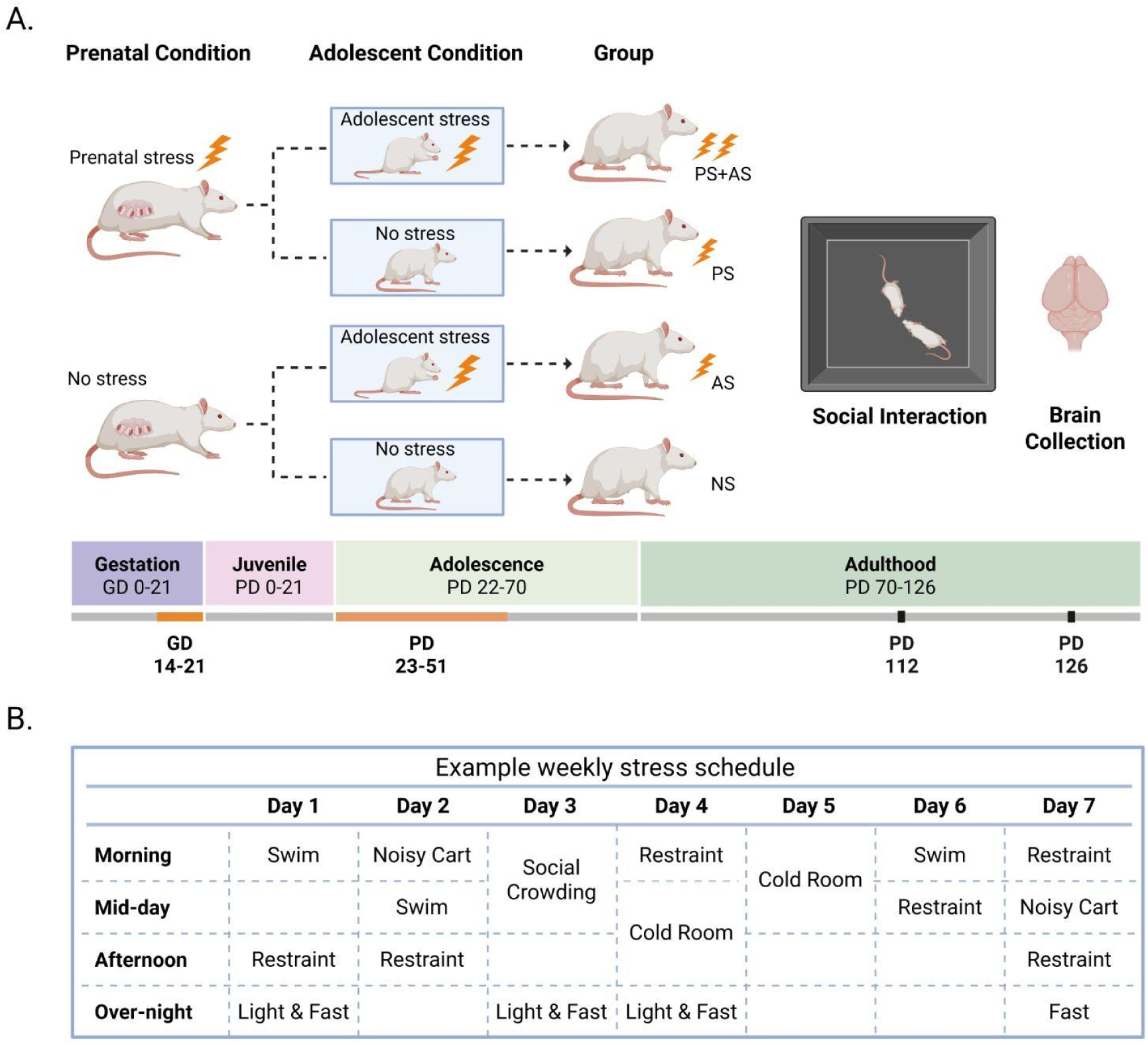
Experimental timeline and an example schedule illustrating chronic unpredictable variable stress procedures. A. Rats were exposed to chronic variable stressors either prenatally (PS) during the last week of gestation (GD 14-21), postnatally during adolescence (AS; PD 23-51), or during both periods combined (PS+AS). A separate group of non-stressed (NS) control animals did not experience stress during either timeframe. Sixty days following stress procedures (PD 112), adult animals underwent an 8-min social interaction test with an unfamiliar sex- and weight-matched non-stressed partner. On PD 126, brains were collected and processed for immunofluorescence of GAD67+ neurons and Wisteria Floribunda Agglutinin to identify PNNs. B. Example 1-week stress schedule. Created using BioRender.com.

### 2.3 Social Interaction Test and Analysis

The social interaction test was performed at 112 days of age in an arena constructed of mat black expanded PVC (70 cm x 70 cm; wall height = 47.6 cm). This test has been previously validated as an ecologically-relevant assay of anxiety-like behavior in rodents (File, 1980; File & Seth, 2003). Experimental subjects interacted with socially-housed partners (2-4/cage) of the same sex that were not subjected to any experimental procedures. The day before testing, all partners were habituated to social testing procedures and received 4 separate 5-min interactions with unfamiliar same-sex animals. Partners were also weighed at this time to permit matching to experimental subjects according to body weight (≤70 g difference between males, ≤ 20 g difference between females). On test day, all animals were transported to the testing suite at 08:00 MDT, and all social interaction tests occurred between 09:00 and 14:00 MDT. The testing room was illuminated by two tall free-standing lamps (71in height; Simple Designs LF2000) placed on opposite sides of the room 3ft equidistant to the testing arena. A plastic shade directed the light (40W) in each lamp toward the ceiling providing indirect illumination. Subjects were allowed 1 minute to explore the open arena before an 8-minute social interaction test began with the introduction of partners. Trials were recorded using ceiling-mounted video cameras, and experimental subjects were ink-marked with an “X” dorsally to facilitate identification during behavioral scoring. The duration of time a subject investigated the face, body, and anogenital region of their partner was scored by a single observer blind to experimental treatment using BORIS event-logging software (Friard & Gamba, 2016). This observer established greater than 90% intra-rater reliability prior to collecting experimental data. Only the sniffing contacts initiated by the experimental subject were scored. Face sniffing was recorded when subjects made sniffing contact with the head, nose, or cheeks of the partner animal. Body sniffing was recorded when the subject made sniffing contact with areas of the partner’s body located between the clavicle and the base of the tail, including the side and flank regions. Anogenital sniffing was scored when the subject made sniffing contact with the area directly underneath the base of the partner’s tail. Sniffing contacts made to the length of the partner’s tail were infrequent and were not scored. Between tests, the arenas were cleaned with Simple Green solution and allowed to dry completely.

### 2.4 Brain collection and sampling

A subset (n = 96) of animals were randomly selected for brain sectioning and histological procedures from animals that underwent social behavior testing (n = 138). This subset resulted in the following number of brains collected per group: PS (n = 24), AS (n = 24), PS+AS (n = 24), PS+AS (n = 24). On postnatal day 126, animals were deeply anesthetized with isoflurane, decapitated, and their brains were rapidly collected and frozen on dry ice before being stored at -80°C. Brains were sectioned coronally at 40 µm using a cryostat (Leica CM1520) maintained at -13 to -12°C. Every third tissue section was mounted onto microscope slides (Superfrost Plus™, 12-550-15), resulting in a sampling interval of 120µm. After sectioning, microscope slides with mounted tissue sections were stored at -80°C until immunofluorescence procedures were conducted to visualize GAD67+ neurons and PNNs.

### 2.5 Immunofluorescence

Slides were thawed at room temperature for 10 minutes and a hydrophobic barrier was drawn around the tissue with a pap pen (VWR, MSPP-SPM0928). Tissue sections were fixed in 4% paraformaldehyde for 10 minutes, followed by three 5-minute washes in phosphate buffered saline (PBS). All subsequent PBS washes described here were also 5 min. After blocking with 10% normal goat serum (G9023, Millipore) for 1 hour, slides were incubated 24 hours at 4°C with mouse anti-GAD67 antibody (Millipore, MAB5406, 1:1000). Slides received three additional PBS washes before a 4-hour incubation at room temperature with goat anti-mouse Cyanine5 (Invitrogen, A10524, 1:1000) and *Wisteria Floribunda Agglutinin* (WFA, Vector, FL-1351-2, 1:500). After two PBS washes, tissue sections were counterstained with DAPI (Invitrogen, D1306, 1:1000) for 5 minutes, washed once more in PBS, and left to dry overnight in the dark. Slides were coverslipped the following day.

### 2.6 Image acquisition and analysis

The BLA complex was identified according to the 6th edition of Paxinos and Watson’s Rat Brain in Stereotaxic Coordinates (Paxinos & Watson, 2007) and spanned bregma -1.56 to -2.76mm. The 120um sampling interval yielded an average of 10 imaged sections per hemisphere. In line with previous studies, the BLA complex was traced as a single region, without subdivision into specific nuclei (Heldt et al., 2012; Pesarico et al., 2022; Smail et al., 2023). Triple immunostained tissue sections were captured bilaterally at 20x magnification (Keyence BZ-X710), using exposure settings optimized to the signal intensity of individual sections as described previously (Lau et al., 2020).

Prior to cell quantification, regional boundaries of the BLA complex were traced using the DAPI channel in ImageJ (Figure 2A; Schneider et al., 2012). Regions were demarcated using visible landmarks from the 6th edition of Paxinos and Watson’s Rat Brain in Stereotaxic Coordinates (Paxinos & Watson, 2007). Two experimenters blind to group assignments established 90% intra- and inter-rater reliability prior to collecting regional mean cross-sectional areas. BLA complex mean cross-sectional areas did not significantly differ between experimental groups (data not presented).

**Figure 2.**
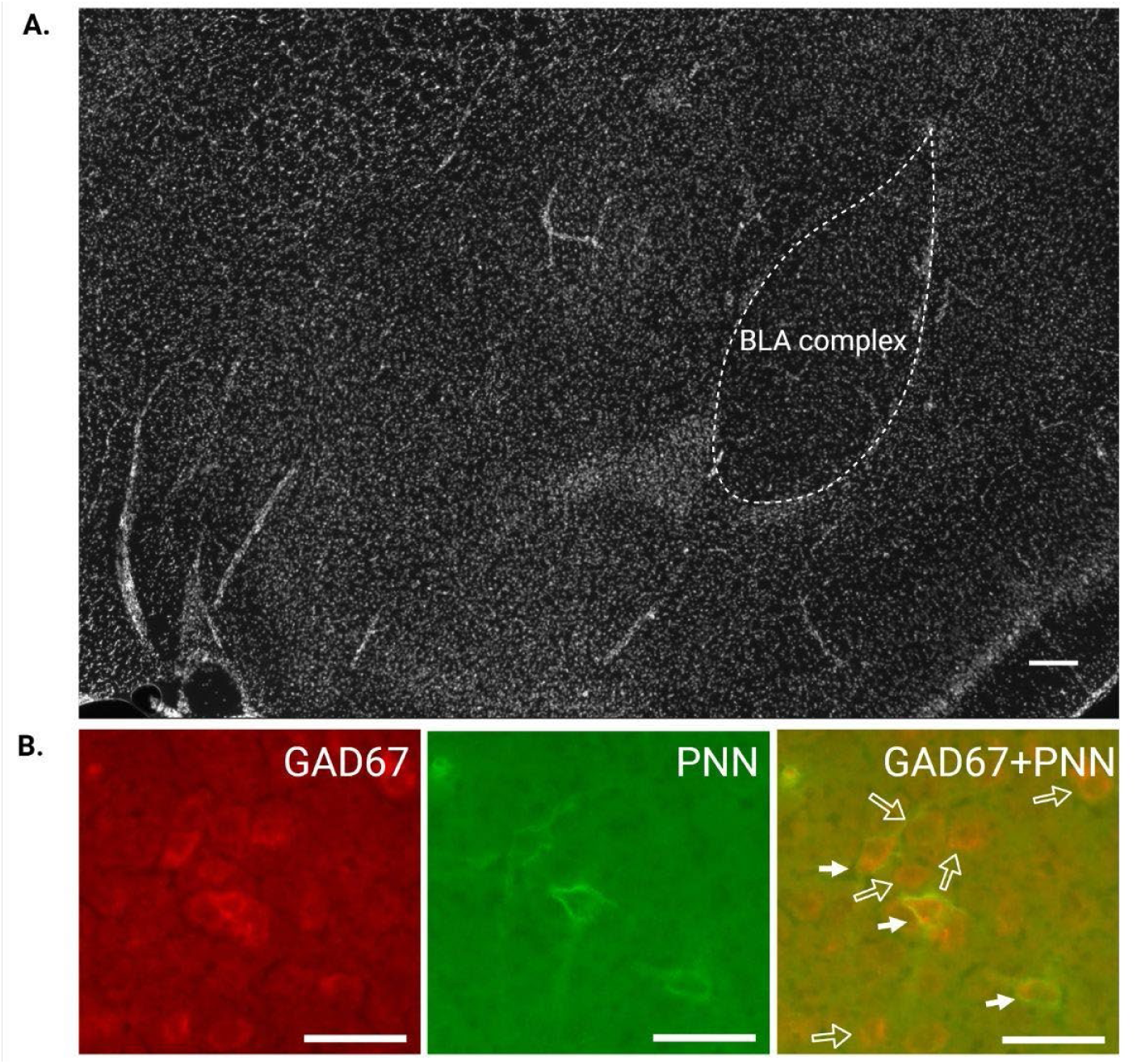
Representative outline of the BLA complex and cell quantification. A. The BLA complex was identified in DAPI prior to GAD67 and PNN quantification. B. GAD67+ neurons were quantified in Cy5, PNNs in GFP, and co-labeled cells were quantified by overlaying Cy5 and GFP. Solid arrows indicate co-labeling of GAD67 and PNN. Open arrows indicate GAD67 with no co-labeling. Scale bar in DAPI = 200 µm; scale bar = 50 µm.

We quantified GAD67+ and WFA+ neurons within each BLA tracing rather than using ROI-based or subsampling methods limited to selected areas within these regions. Specific morphological and labeling criteria were used to quantify GAD67+ neurons and PNNs. GAD67+ neurons exhibited intense fluorescence in the cytoplasm of the soma but not within the nucleus (Figure 2B). PNNs were quantified if WFA labeling encompassed at least 80% of a cell soma and co-labeled cells were quantified using the multipoint tool in ImageJ. The number of GAD67+ neurons, PNNs, and number of PNNs surrounding GAD67+ neurons (i.e., co-labeled cells) were quantified. Two individuals blind to experimental conditions achieved intra- and inter-rater reliability of at least 90% before quantifying the number of GAD67+ neurons, PNNs, and the number of co-labeled cells on unmodified images.

### 2.7 Experimental Attrition and Data analysis

For behavioral analyses, the data from two female and two male subjects were unusable due to a noise disruption during their behavioral tests. For BLA analyses, data from four animals were not analyzed due to loss of tissue during histological processing. Final sample sizes for experimental groups for the BLA were as follows: NS (n = 23), PS (n = 22), AS (n = 23), PS+AS (n = 24). Three-factor between-subjects ANOVAs (sex x prenatal condition x adolescent condition) were conducted to analyze the total time subjects spent actively investigating the partner, and the time subjects spent sniffing different areas of the partner (body, face, and anogenital region). The BLA was analyzed by mixed ANOVA treating sex, prenatal condition, and adolescent condition as between-subject factors, and brain hemisphere as the within-subjects repeated measure. The dependent measures included the number of GAD67+ neurons, WFA+ neurons, the number of cells co-labeled with GAD67 and WFA, and the percentage of GAD67+ neurons co-labeled with WFA. In addition to p-values, measures of effect size (partial eta squared, ηp²) are also reported.

## 3. Results

### 3.1 Prenatal stress exposure prevents adolescent stress-induced increases in social investigation

Prenatal and adolescent conditions interacted to impact subject social investigation duration (Figure 3A; *F*_(1,133)_ = 9.58, *p* = .002, ηp² = .071). Adolescent stress significantly increased the time subjects investigated their partner relative to NS controls (p<.001). AS subjects also investigated partners for significantly greater durations than did subjects exposed to both prenatal and adolescent stress (p<.001), suggesting that prenatal stress exposure prevented the impact of adolescent stress on social investigation in PS+AS subjects (Figure 3A). Male and female subjects did not differ in time spent investigating partners (*F*_(1,133)_ = .000, *p* = .994, ηp² = .000), nor did sex interact with prenatal or adolescent condition. However, interesting effects of sex emerged when the time subjects spent investigating their partner’s body, face, and anogenital region was examined. Both male and female subjects spent the majority of test time sniffing the body of their partners, but males exhibited overall greater durations than females (Figure 3B; *F*_(1,133)_ = 10.45, *p* < .01, ηp² = .077). Although main effects of prenatal (*F*_(1,133)_ = 8.42, *p* < .01, ηp² = .063) and adolescent condition (*F*_(1,133)_ = 4.97, *p* < .03, ηp² = .038) on body sniffing were also observed, these factors did not interact with sex given that a similar pattern of group differences was observed in both males and females (Figure 3B). Unique effects of stress were observed between males and females for face sniffing duration. Although main effects of sex (p < .001), prenatal (p < .05), and adolescent condition (p < .02) were found, these effects were qualified by a significant 3-way interaction (Figure 3C; *F*_(1,133)_ = 4.47, *p* < .04, ηp² = .034). In males, face sniffing duration followed a similar pattern to that observed for the overall social investigation duration. Specifically, adolescent stress significantly increased the duration of face sniffing relative to nonstressed controls (p < .001), and AS subjects also engaged in longer face sniffing durations than did PS+AS subjects (p < .001), suggesting that prenatal stress prevents adolescent stress-induced increases in face sniffing in PS+AS males (Figure 3C). In females, no effects of prenatal or adolescent condition on face sniffing durations were observed. For anogenital investigation, females spent overall longer durations than males investigating this body region of their partner (Figure 3D; *F*_(1,133)_ = 52.25, *p* < .001, ηp² = .293). Sex also interacted with adolescent condition (*F*_(1,133)_ = 13.30, *p* < .001, ηp² = .096), and with both prenatal and adolescent condition (*F*_(1,133)_ = 3.60, *p* = .06, ηp² = .027). In females, adolescent stress significantly increased anogenital investigation relative to nonstressed controls (p < .001). AS females also sniffed the anogenital region for significantly longer durations than did PS+AS females (p < .001), indicating that prenatal stress exposure changed the impact of adolescent stress on anogenital investigation (Figure 3D). In males, no effects of prenatal condition, adolescent condition, or interactions between these factors were detected for anogenital investigation. Taken together, although the total investigation duration (Figure 3A) did not differ between males and females, adolescent stress induced unique targets for social investigation: the anogenital region in females, and the face in males.

**Figure 3.**
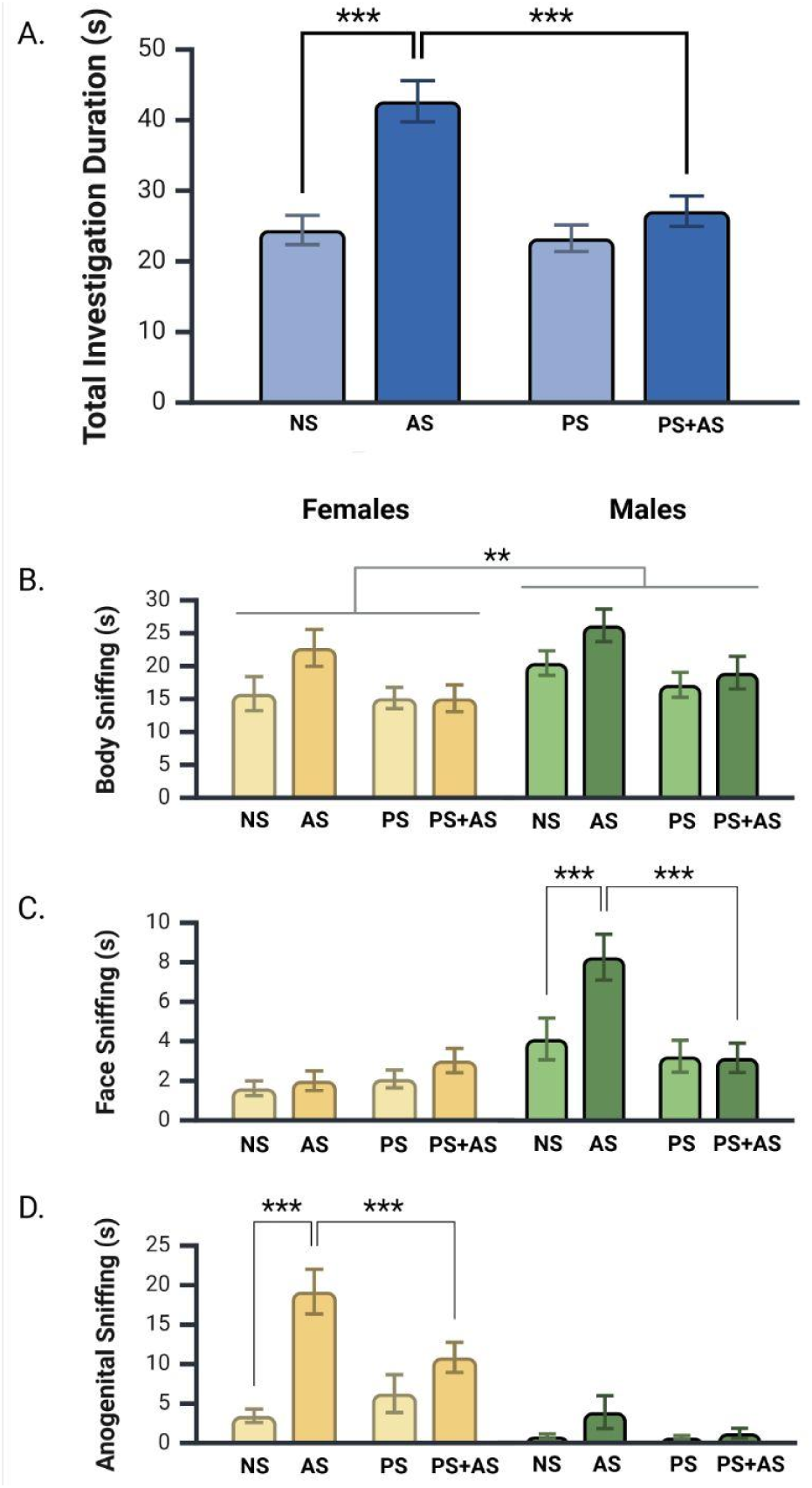
The effects of prenatal (PS), adolescent (AS), and combined prenatal and adolescent stress (PS+AS) on social investigation of a same-sex and weight-matched partner. A. Adolescent stress significantly increased social investigation of a partner relative to nonstressed (NS) controls. Adolescent stressed subjects also displayed longer investigation durations than subjects than PS+AS subjects, suggesting that prenatal stress exposure changes the effects of adolescent stress on investigatory behavior. B. Males investigate the body region of partners for longer durations than females, overall. Males and females exhibit a similar stress-dependent pattern of body sniffing C. Adolescent stress increases face sniffing durations in male but not female subjects. The pattern of group differences in male face sniffing is similar to overall investigation duration. D. Adolescent stress increases anogenital sniffing durations in female but not male subjects. A-D. Darker shading indicates the presence of adolescent stress, and lighter shading indicates the absence of adolescent stress. Bars represent group mean +/- SEM. * indicates p < .05, ** indicates p < .01, and *** indications p < .001.

### 3.2 Prenatal stress exposure prevents adolescent stress-induced decreases in the number and percentage of GAD67+ neurons surrounded by PNNs in the BLA complex

The number of GAD67+ neurons was not impacted by sex (*F*_(1,84)_ = 003, *p* = .95, ηp² =.000) or hemisphere (*F*_(1,84)_ = .242, *p* = .62, ηp² = .003), therefore, the data are collapsed across sex and hemisphere in Figure 4A. Neither prenatal condition (*F*_(1,84)_ = 1.00, *p* = .319, ηp² = .012) nor adolescent condition (Figure 4A; *F*_(1,84)_ = .128, *p* = .72, ηp² =.002) affected GAD67+ neuron number. Similarly, PNN number was not significantly affected by sex, (Figure 4B; *F*_(1,84)_ = .093, *p* = .761, ηp² = .001), hemisphere, (*F*_(1,84)_ = .016, *p* = .901, ηp² = .000), prenatal condition (*F*_(1,84)_ = .224, *p* = .637, ηp² = .003), or adolescent condition, (*F*_(1,84)_ = .162, *p* = .688, ηp² = .002). Thus, if prenatal or adolescent stress altered the number of GAD67+ neurons or PNNs, these effects did not persist into adulthood.

**Figure 4.**
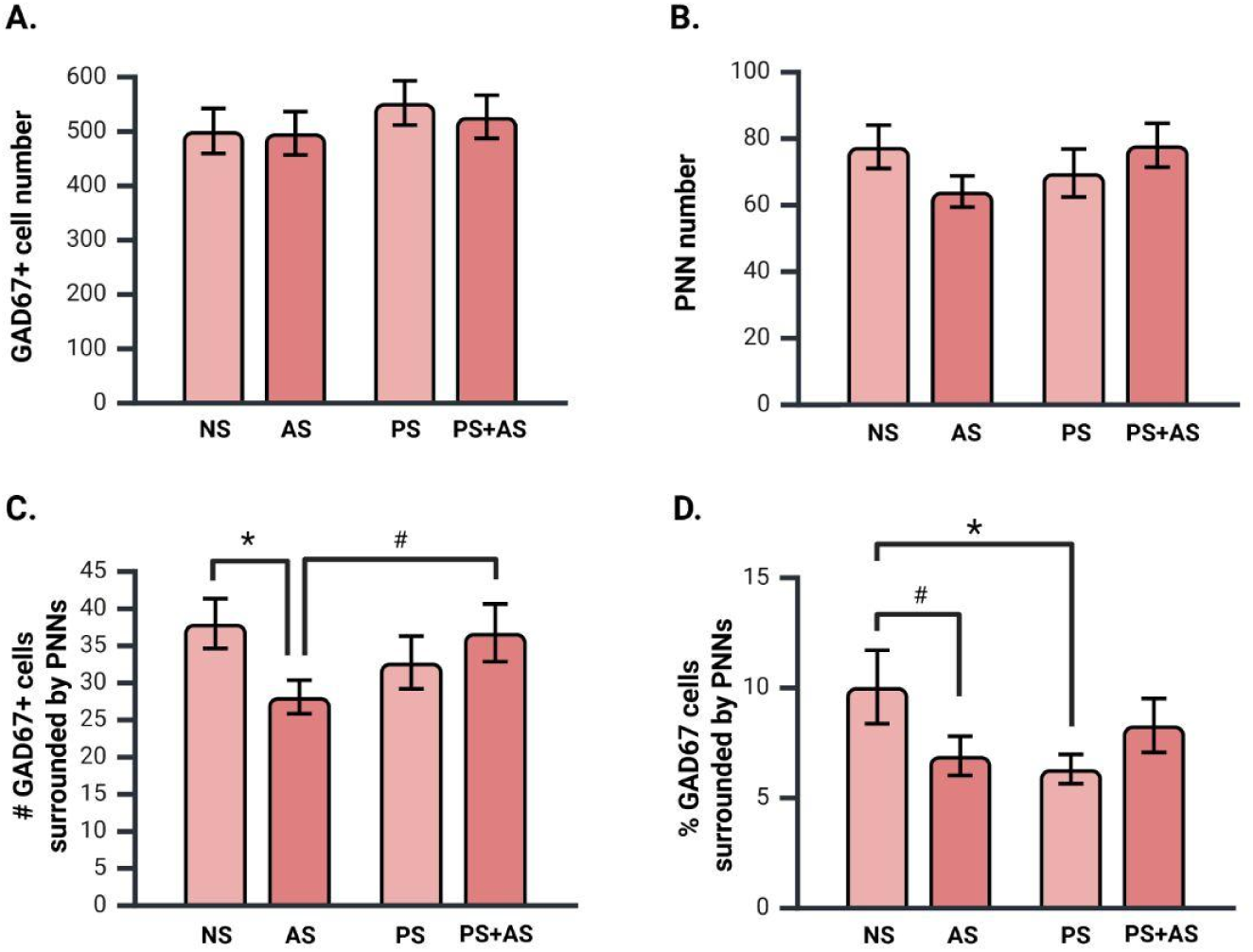
The effects of prenatal (PS), adolescent (AS), and combined prenatal and adolescent stress (PS+AS) on inhibitory neurons in the BLA. A. Stress did not influence GAD67+ neuron number. B. The number of PNNs was not affected by prenatal or adolescent stress. C. Relative to nonstress (NS) controls, AS significantly decreased the number of GAD67+ cells surrounded by PNNs (co-labeled). In contrast, combined PS+AS co-labeling did not differ from control, and was greater than AS alone. This data pattern suggests that when preceded by prenatal stress, adolescent stress no longer decreases the number of PNNs surrounding inhibitory neurons. D. Compared to NS controls, both AS and PS decreased the percentage of GAD67+ neurons surrounded by PNNs, whereas combined PS+AS did not differ from control. A-D. Dark red bars indicate the presence of adolescent stress, and light red bars indicate the absence of adolescent stress. Average left and right hemisphere values are presented, and bars represent group mean +/- SEM. * indicates p < .05, # indicates p ≤ .07.

Prenatal and adolescent conditions interacted to influence the number of GAD67+ neurons surrounded by PNNs (*F*_(1,84)_ = 4.127, *p* = .045, ηp² = .047). AS reduced the number of GAD67+ neurons surrounded by PNNs compared to non-stressed controls (p = .04) and animals that experienced combined prenatal and adolescent stress (p = .07). Additionally, PS+AS animals did not significantly differ from nonstressed controls (p =.789). Overall, this data pattern suggests that when stressors are combined, prenatal stress exposure prevents adolescent stress-induced reductions in the number of GAD67+ neurons surrounded by PNNs (Figure 4C). The number of GAD67+ neurons co-labeled with PNNs was not influenced by sex, (Figure 4C; *F*_(1,84)_ = .014, *p* = .907, ηp² = .000), or hemisphere, (*F*_(1,84)_ = .007, *p* = .935, ηp² = .000), therefore the data are collapsed across sex and hemisphere in Figure 4C.

Prenatal and adolescent condition also interacted to influence the percentage of GAD67+ neurons surrounded by PNNs (Figure 4D; *F*_(1,84)_ = 4.658, p = .03, ηp² = .053). Compared to non-stressed animals, a reduction in the percentage of GAD67+ neurons surrounded by PNNs was observed in both prenatal stress (p = .03) and adolescent stress (*p =* .06) groups. However, no difference was observed between the combined PS+AS group and non-stressed controls (p = .47). The percentage of GAD67+ neurons surrounded by PNNs was not dependent upon on sex (*F*_(1,84)_ = .191, *p* = .66, ηp² = .002) or hemisphere (*F*_(1,84)_ = .030, *p* = .86, ηp² = .000), therefore, the data are collapsed across sex and hemisphere in Figure 4D.

### 3.3 Terminal Body Weights

Table 2 provides mean terminal body weights for each group. As expected, body weights were significantly greater in male than in female subjects (*F*_(1,84)_ = 789.72, *p* <.001, ηp² = .904). A marginally significant main effect of adolescent condition was present (*F*_(1,84)_ = 3.5, *p* = .06, ηp² = .040), indicating that animals exposed to adolescent stress (AS, PS+AS) tended to weigh less than those not exposed to adolescent stress (NS, PS), irrespective of sex or prenatal condition. No interactions between sex, prenatal, or adolescent condition were observed.

**Table 2.** Mean (SD) terminal body weights.

|  | Group | Weight (g) |
| --- | --- | --- |
| Females | NS | 319.28 (36.74) |
|  | AS | 293.83 (26.76) |
|  | PS | 321.00 (31.46) |
|  | PS+AS | 294.50 (25.48) |
| Males | NS | 587.23 (67.12) |
|  | AS | 576.75 (48.24) |
|  | PS | 573.90 (58.32) |
|  | PS+AS | 565.25 (49.96) |

## 4. Discussion

The current study sought to determine whether the effects of stress experienced during both prenatal and adolescent development are additive, or whether prenatal stress changes the impact of adolescent stress. We provide here the first evidence that prenatal stress exposure prevents adolescent stress-induced *decreases* in the number and percentage of BLA inhibitory neurons surrounded by PNNs, and also prevents adolescent stress-induced *increases* in adult social investigation. These findings align with the predictive-adaptive-response (PAR) model proposing that prenatal experiences serve as cues to predict the postnatal environment and stimulate adaptive responses that “match” the phenotype to this prediction (Gluckman et al., 2005; Gluckman & Hanson, 2004).

Few studies to date have investigated interactions between prenatal and adolescent stress exposures on adult brain and behavioral outcomes. Lee et al., (2016) demonstrated that although stressed male mice exhibit impaired spatial memory function in adulthood, no adult memory impairments are observed in males exposed to combined prenatal and adolescent stress (Y.A. Lee et al., 2016). While our study investigated the impact of stress exposure during two developmental periods (prenatal and adolescence), the majority of studies have employed designs in which stress exposures occur during a single developmental period (e.g. prenatal, early postnatal, or adolescent periods) and adulthood. For example, prenatally stressed male rats cope more effectively in adulthood with the stress of a subordinate social status than do subordinates that were not exposed to prenatal stress (Scott et al., 2017). In mice, restricting nesting and bedding materials during the juvenile period mitigates the anxiogenic effects of chronic social defeat and social isolation in adulthood (Santarelli et al., 2014, 2017). Similarly, rats experiencing CUS in adolescence are resistant to the detrimental effects of adult stress exposure on emotional memory function (Chaby et al., 2020). Collectively, these studies demonstrate that the effects of adult stress on behavior depend on stress history during postnatal development, consistent with our findings demonstrating that the effects of adolescent stress on adult social behavior depend on prenatal stress history.

We report here that stress isolated to the adolescent period caused a robust increase in social investigation in adulthood, a finding that aligns with several previous studies employing social isolation stress during adolescence (Lander et al., 2017; Manojlović et al., 2024; Rivera-Irizarry et al., 2020); Burke et al., 2025; Dawud et al., 2021; Fontenot et al., 2018; Goodell et al., 2017; Wall et al., 2012). To our knowledge, we are the first to report that exposure to CVS throughout adolescence causes long-term increases in social investigation in both male and female rats. Our findings contrast with two previous studies in male Sprague Dawley rats that observed *decreases* in social behavior following adolescent stress using similar CVS procedures (Oztan et al., 2011; Toth et al., 2008). However, because social interaction testing occurred *immediately* following the cessation of adolescent CVS procedures (Oztan et al., 2011; Toth et al., 2008), the acute effects of CVS on social behavior were indistinguishable from the potential long-term impacts of adolescent stress exposure in these previous studies. While our data suggest that in the long-term, adolescent exposure to CVS increases social investigation in both male and female Sprague Dawley rats, additional studies are needed to determine whether the effects of adolescent CVS exposure change across time and/or are dependent upon subject age at behavioral testing. Thus, while numerous studies report lasting effects of adolescent stress on social behavior, whether adolescent stress-induced increases or decreases in social behavior are observed likely depends on the specific stress protocols implemented, the timing of stress exposure within the adolescent period, and the timing of behavioral testing relative to the cessation of stress procedures (McCormick, 2021).

The expression of social behavior in rats is sensitive to environmental context, and social interactions decrease between adult rats when the testing environment is unfamiliar (File, 1980; File & Seth, 2003; Varlinskaya & Spear, 2008). Although we did not test animals in both familiar and unfamiliar test contests, in an unfamiliar testing chamber, subjects exposed to adolescent stress alone displayed higher levels of social investigation than all other groups. This data pattern suggests that adolescent stress may disrupt the context-appropriate expression of social behavior in adulthood. Adolescent stress also caused sex-specific changes in investigation preferences. AS females preferentially investigated the anogenital region of partners, whereas AS males preferentially investigated their partner’s face. Dominance status is bidirectionally communicated between male rats during face to face encounters, and a decrease in the subordinate male’s sniffing frequency signals their submissive status and prevents an escalation in aggression (Wesson, 2013). Thus, it is possible that increased face sniffing in AS males reflects an increased motivation to dominate a social partner. In line with this possibility, adolescent social isolation stress increases the motivation of male rats to interact with an aggressive partner in late adolescence (Goodell et al., 2017).

In adult rodents, decreased inhibition and/or increased excitation of the BLA decreases social interactions (Asim et al., 2024; Paine et al., 2017; Rosenkranz et al., 2010; Sanders & Shekhar, 1995; Truitt et al., 2007), and previous studies indicate that stress-induced BLA excitation is the consequence of decreased inhibitory regulation (Asim et al., 2024; Truitt et al., 2007). In the current study, adolescent stress decreased the number of GABAergic neurons surrounded by PNNs, however, the influence of PNNs on the excitatory/inhibitory balance of the BLA is not yet known. In brain regions such as the mPFC and visual cortex, removal of PNNs decreases inhibitory function (Carceller et al., 2020; Lensjø, Christensen, et al., 2017). Thus, additional studies are required to determine whether the observed adolescent stress-induced decreases in inhibitory neurons surrounded by PNNs contributes to changes in BLA excitatory/inhibitory balance and levels of social investigation in adulthood.

Our findings in the BLA complement and extend previous studies investigating the effects of stress exposure during adolescence on PNN development within cortico-amygdala neural circuits that regulate social and anxiety-related behavior. Few studies have investigated the long-term effects of adolescent stress on PNNs (Colodete et al., 2024; Page & Coutellier, 2018; Santos-Silva et al., 2024). We report here that adolescent stress reduces the number and percent of inhibitory neurons surrounded by PNNs in the adult BLA, a finding that aligns with other reports that adolescent stress decreases the number of parvalbumin (PV)-containing interneurons surrounded by PNNs in the medial prefrontal cortex (Santos-Silva et al., 2024) and ventral hippocampus in adulthood (Colodete et al., 2024). To our knowledge, we provide here the first evidence that similar long-term effects of adolescent stress also occur in the BLA. Our study differs from others, however, in that we quantified all inhibitory neurons rather than only those containing PV. The BLA is composed of many inhibitory neuronal subtypes (Vereczki et al., 2021), and the changes we observed in the number and percentage of inhibitory neurons with PNNs may reflect stress-induced alterations in PV, as reported in other studies. Overall, our findings add and extend to limited but growing evidence that the effects of developmental stress persists into adulthood and leads to long-lasting alterations in inhibitory neurons and PNNs within the BLA.

Stress-dependent alterations in microglia function may underlie the stress history-dependent effects of adolescent stress on PNN-surrounded inhibitory neurons reported here. Microglia, the resident immune cells of the central nervous system, disassemble PNNs when in an activated pro-inflammatory state (Crapser et al., 2020; Liu et al., 2024; Tansley et al., 2022; Venturino et al., 2021). Chronic stress is a well-established trigger of microglial activation (for review Schramm & Waisman, 2022). and early-life stress can alter microglial responsiveness to future stressors. For example, adolescent stress exposure increases microglia reactivity in adulthood (Rodríguez-Arias et al., 2018; Wang et al., 2018). While speculative, the adolescent stress-dependent decrease in PNN-surrounded inhibitory neurons we report here may be the result of adolescent stress-induced microglia activation and disassembly of PNNs. In contrast, other studies indicate that prenatal immune stress can dampen microglial responses to later immune challenges (Hayes et al., 2022; Schaafsma et al., 2017). Given this, the effects of repeated stress on microglia function may also explain why adolescent stress failed to decrease PNNs when preceded by prenatal stress. Specifically, when animals experienced combined PS+AS, prenatal stress exposure may have blunted microglia reactivity and disassembly of PNNs in response to subsequent adolescent stress. Despite the abundance of studies demonstrating the effects of developmental stress exposure on both microglia and PNNs, and that demonstrate microglia-dependent disassembly of PNNs, experiments directly linking stress exposure with microglia-dependent changes in PNNs are lacking. Studies are currently underway in our laboratory that directly test whether microglia activation mediates the effects of prenatal and adolescent stress on PNNs within the BLA and other stress-sensitive brain regions.

Overall, our data demonstrate that prenatal stress prevents the long-lasting effects of adolescent stress on inhibitory neurons and PNNs in the BLA. When experienced in isolation, adolescent stress reduced the number and percentage of GAD67+ neurons surrounded by PNNs, and increased social investigation. However, when prenatal stress preceded adolescent stress exposure, the effects of combined PS+AS were indistinguishable from nonstressed controls. As such, prenatal stress exposure may confer resilience to the effects of adolescent stress on inhibitory neurons in the BLA and normalize investigatory behavior during social interactions.

## 5. CRediT authorship contribution statement

**Marcia Chavez**: Methodology, investigation, project administration, data curation, formal analysis, visualization, writing – original draft, writing – review & editing. **Billy Lau**: Methodology, writing – review & editing. **Madeline Jones:** Investigation, writing – review & editing. **Alyssa Whaley** Investigation, writing – review & editing. **Trevor Pounders**: Investigation, writing – review & editing. **Kalynn Schulz**: Conceptualization, funding acquisition, resources, methodology, investigation, supervision, data curation, visualization, writing – review & editing.

## 8. Acknowledgements

We thank Ansley Inscore and Isabella Rochford for their assistance with brain imaging, and Krystiana Rego for assistance with data entry. We gratefully acknowledge Arthur Jay Castaneda, Sierra Lawler, and Krystiana Rego for providing valuable feedback on this manuscript.

